# Cell-type and region-specific nucleus accumbens AMPAR plasticity associated with morphine reward, reinstatement, and spontaneous withdrawal

**DOI:** 10.1101/592519

**Authors:** Aric C. Madayag, Devan Gomez, Eden M. Anderson, Anna E. Ingebretson, Mark J. Thomas, Matthew Hearing

**Affiliations:** Department of Biomedical Sciences, Marquette University, Milwaukee, WI, 53233; Department of Neuroscience, University of Minnesota, Minneapolis, MN 55455

**Author notes:** **Corresponding Author:** Matthew C. Hearing, PhD.

## Abstract

Despite evidence that morphine-related pathologies reflect adaptations in NAc glutamate signaling, substantial gaps in basic information remain. The current study examines the impact of non-contingent acute, repeated, and withdrawal-inducing morphine dosing regimens on glutamate transmission in D1- or D2-MSNs in the NAcSh and NAcC sub-regions in hopes of identifying excitatory plasticity that may contribute to differing facets of opioid addiction-related behavior. Three hours following an acute morphine injection (10 mg/kg), average miniature excitatory postsynaptic current (mEPSC) amplitude mediated by AMPA-type glutamate receptors was increased at D1-MSNs in the both the shell and core regions, whereas only the frequency of events was elevated at D2-MSNs in the shell. In contrast, somatic withdrawal induced by escalating dose of repeated morphine twice per day (20, 40, 60, 80, 100mg/kg) only enhanced mEPSC frequency at D2-MSNs in the shell 24 hrs following the final drug exposure. Further, drug re-exposure 10-14 days following a preference-inducing regimen of morphine produced a rapid and enduring endocytosis of GluA2-containing AMPARs at D1-MSNs in the shell, that when blocked by an intra-NAc shell infusion of the Tat-GluA2_3Y_ peptide, increased reinstatement of morphine place preference – a phenomenon distinctly different than effects previously found with cocaine. The present study is the first to directly identify unique circuit specific adaptations in NAc glutamate synaptic transmission associated with morphine-related acute reward and somatic withdrawal as well as post-abstinence short-term plasticity. While differing classes of abused drugs (i.e., psychostimulants and opioids) produce seemingly similar bidirectional plasticity in the NAc following exposure to relapse-linked stimuli, our findings indicate this plasticity has distinct behavioral consequences.

**Compliance with Ethical Standards:** The authors have no conflicts of interest to disclose. All authors have given their consent for manuscript submission. The research in the current study used mice single- or group-housed on a 12 h light/dark cycle with food and water available ad libitum with experiments run during the light portion. All experiments were approved by the University of Minnesota and Marquette University Institutional Animal Care and Use Committee. The following funding sources made the study possible: National Institute for Neurological Disorders and Stroke (P30 NS062158); National Institute on Drug Abuse grant K99 DA038706 (to M.H.), R00DA038706 (M.H.), R00DA038706-04S1 (A.C.M), R01DA019666 (M.J.T.), K02DA035459 (M.J.T.) and T32 DA007234 (A.E.I.).

## INTRODUCTION

Opioids are the main class of drugs for pain management despite the risk for abuse (Wise 1989). In addition to their primary rewarding effects, repeated use of opioids can result in the development of physical dependence that manifests as debilitating somatic and psychological withdrawal symptoms that can perpetuate continued use (Koob et al. 1989; van Ree et al. 1999). Increasing evidence suggests that opioid-induced plasticity related to tolerance, dependence, and withdrawal occurs within divergent, as well as overlapping, neural circuits as plasticity responsible for establishing opioid-seeking behavior and drug-associated stimuli that can provoke craving and relapse (Badiani et al. 2011; Graziane et al. 2016; Hearing 2019; Hearing et al. 2018; Hearing et al. 2016; Russell et al. 2016; Zhu et al. 2016) - highlighting a major challenge towards identifying the neurophysiological bases of dependence and withdrawal versus adaptations responsible for enduring relapse risk.

Prior findings posit glutamate plasticity in the nucleus accumbens (NAc) as a significant factor in the acute rewarding effects of opioids (Baharlouei et al. 2015), conditioned opioid-associations (Fujio et al. 2005; Hearing et al. 2016; Siahposht-Khachaki et al. 2017), and relapse vulnerability (Bossert et al. 2005; Bossert et al. 2006; Shen et al. 2011; Shen et al. 2014). Data also indicate that elevations in NAc glutamate transmission underlie somatic and affective withdrawal symptoms (Russell et al. 2016; Sepulveda et al. 2004; Zhu et al. 2016). However, the NAc is a heterogeneous area of the brain divided into NAc core (NAcC) and shell (NAcSh) subregions based on anatomical connectivity. While the NAcC subregion interacts with brain regions associated with motor circuitry, thus coordinating behavioral output, the NAcSh interacts with limbic and autonomic brain regions, indicating significant regulation of reward, emotional, and visceral responses to stimuli (Everitt et al. 1999; Heimer et al. 1991; Zahm and Brog 1992). Within each subregion, the primary target of excitatory glutamate afferents are the principal medium spiny projection neurons (MSNs), which are categorically divided based on expression of type 1 (D1-MSNs) or type 2 dopamine receptors (D2-MSNs) (Le Moine and Bloch 1995; Lobo and Kennedy 2006; Smith et al. 2013).

Despite evidence that morphine-related pathologies reflect adaptations in NAc glutamate signaling, substantial gaps in basic information remain. For example, while acute morphine exposure transiently increases extracellular glutamate in the NAc (Desole et al. 1996; Enrico et al. 1998; Sepulveda et al. 2004), evidence supporting a role of AMPAR plasticity is lacking. Further, while elevations in NAc shell GluA1-containing AMPA-type receptors has recently been shown to causally contribute to morphine dependence increases expression of GluA1-containing AMPA-type receptors (Russell et al. 2016), it remains unclear whether similar changes occur during spontaneous withdrawal, and in what cell-type these adaptations occur. Increasing evidence indicates the nature and locus of opioid-induced glutamate plasticity in the NAc dictates the relationship to behavior, with most findings to date highlighting adaptations to the NAcSh in opioid reward and aversion (Gracy et al. 2001; Graziane et al. 2016; Hearing et al. 2016; Russell et al. 2016; Svingos et al. 1997; Zhu et al. 2016). For example, abstinence from non-contingent morphine administration is associated with divergent plasticity in the NAcSh at D1- and D2-MSNs (Graziane et al. 2016; Hearing et al. 2016), but not in the NAcC (Hearing et al. 2016), with increased transmission at D1- and D2-MSNs contributing to opioid reward and aversion learning, respectively (Graziane et al. 2016; Hearing et al. 2016; Russell et al. 2016; Zhu et al. 2016). The current study examines the impact of non-contingent acute, repeated, and withdrawal-inducing morphine dosing regimens on glutamate transmission in D1- or D2-MSNs in the NAcSh and NAcC sub-regions in hopes of identifying excitatory plasticity that may contribute to differing facets of opioid addiction-related behavior.

## Materials and Methods

### Animals

Adult (P49-72) male mice were a combination of heterozygous bacterial artificial chromosome (BAC) transgenic mice (Jackson Laboratories, Bar Harbor, ME, USA) expressing tdtomato or eGFP expression driven by either dopamine receptor DR1 (drd1a-tdtomato) or DR2 (drd2-eGFP), or double transgenics expressing tdtomato and eGFP. Mice were single- or group-housed on a 12 hrs light/dark cycle with food and water available ad libitum with experiments run during the light portion. All experiments were approved by the University of Minnesota and Marquette University Institutional Animal Care and Use Committees.

### Stereotaxic intra-cranial cannula implantation

For surgical procedures, mice were anesthetized with ketamine and xylazine (100/10 mg/kg, respectively, i.p.). Depth of anesthesia was assessed prior to the subject being placed in the stereotaxic frame (Kopf Instruments, Tujunga, CA, USA). Measurements targeting implantation of the single barrel guide cannula (26ga, 5mm pedestal, 3.5mm projection; C315GS-5/SP, Plastics1, Roanoke, VA, USA) to the NAcSh region were taken with respect to bregma/midline (+1.50 a/p, +/− 1.45 m/l, −4.0) at a 14° angle. Cannula were cemented in place using Geristore (DenMat, Lompoc, CA, USA). Mice were allowed a minimum 5-day recovery period before beginning behavior testing.

### Morphine-induced Locomotion

Locomotor chamber apparatus was placed under AnyMaze video tracking system (Stoelting, Wood Dale, IL, USA) and measurements were made automatically by the software as previously described (Hearing et al. 2016).

### Acute Morphine

Mice were given an injection (i.p.) of saline or morphine (10 mg/kg) and euthanized 3-4 hrs following injection for electrophysiological recordings. This time point was chosen for the purpose of performing the electrophysiological recordings while morphine was still present in tissue and serum. Our recordings were performed prior to reaching the approximate 5-hour half-life of subcutaneous morphine administration (Hipps et al. 1976) and during a post-injection time point previously shown to observe elevated locomotor activity (Hearing et al. 2016). These experiments were performed at the University of Minnesota.

### Morphine Challenge

Mice were administered 5 daily injections of saline or morphine (10 mg/kg) – a regimen previously shown to augment glutamate transmission and promote sensitization (Hearing et al. 2016) – followed by 10-14 days of abstinence. Mice were then administered a saline or morphine (10 mg/kg) challenge injection. 16-24 hrs following challenge injection, mice were euthanized for subsequent electrophysiological recordings. These experiments were performed at both the University of Minnesota and Marquette University.

### Spontaneous Withdrawal

Across a period of 5 days, 2 injections (saline or morphine) were given each day in their home cage approximately 12 hrs apart, with doses for each day at 20, 40, 60, 80, and 100 mg/kg (10 injections total). Twenty-four hours or 14 days following the final injection, mice were placed into a clear plastic cage (16.5”x8”x7.5”) and examined for signs of somatic withdrawal during a 30 min period. Somatic measures were chosen based on previous works examining morphine withdrawal (Cruz et al. 2008; Papaleo and Contarino 2006; Schulteis et al. 1994; van der Laan and de Groot 1988; van der Laan et al. 1991). Jumps, tremors, paw flutters, wet dog shakes, piloerection, and grooming were hand scored with each measurement recorded as a score of one, including the singular possible observation of piloerection, to generate a global withdrawal score with the following equation: 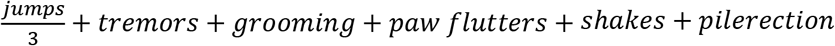 (Papaleo and Contarino 2006). Additional measurements included locomotor activity in the form of distance traveled (meters, m) using AnyMaze video tracking software (Stoelting Company, Wood Dale, IL, USA). These experiments were performed at Marquette University.

### Conditioned Place Preference

All conditioned place preference experiments employed a two-chamber apparatus (St. Albans, VT, USA) and were performed as previously described (Wydeven et al. 2014). For conditioning, subjects were injected with morphine (5 mg/kg) or vehicle, and after a 20-min delay were confined for 30 min in the corresponding CS+/CS-chamber. Morphine dosing for place preference training was chosen based on previous findings that this dose produces robust place preference (Hearing et al., 2016). Further, this dose in addition to extinction training following conditioning was shown to produce identical cell-specific plasticity observed following 5 daily injections of morphine and home cage abstinence (Hearing et al. 2016). A total of four morphine (5 mg/kg) and four saline trials were performed in alternating fashion, with only one trial performed per day and preference evaluated 24 hrs following the final conditioning session. Following conditioning, mice underwent 6 daily extinction sessions as previously described (Hearing et al. 2016), with animals confined to the CS+ and CS-compartment for 20 min each on days 1, 3, and 5 (extinction training) and allowed to freely explore on days 2, 4, and 6 (extinction testing). Day 6 data was used for two-way ANOVA analysis.

### Intra-cranial GluA2 peptide and reinstatement of place preference

Experimental treatments for the reinstatement tests were assigned after extinction training and assignments were made to ensure that each treatment group had similar preference scores prior to and following extinction. Reinstatement of place preference was performed in five different experiments. To block endocytosis of GluA2-containing AMPA receptors in the NAcSh, a synthetic interference peptide designed to disrupt activity-dependent endocytosis without altering basal receptor trafficking, was used as previously described (Ahmadian et al. 2004; Brebner et al. 2005). Mice received an intra-NAcSh infusion of the active (Tat-GluA2_3Y_) or inactive version (Tat-GluA2_3A_) of the peptide diluted in ACSF (75 pmol; 0.5 μL/hemisphere; 0.1 μL/min) using a 32ga internal cannula with 1.2 mm projection beyond the guide. Following infusions, mice were returned to their home cage for 60 min, at which point they received an i.p. injection of morphine (5 mg/kg) or saline, followed by examination of preference behavior during a 20 min test. Electrophysiology recordings to confirm effects of Tat-peptide expression were done within 2 hrs following testing. These experiments were performed at the University of Minnesota.

### Histological Analysis

Histological examination of cannula targeting was done visually on the electrophysiology rig or post-mortem in tissue fixed with transcardial perfusion of 4% paraformaldehyde buffered in saline using an overdose of pentobarbital (650 mg/kg). Brains were cryoprotected, sliced at 40μm, mounted, and cover-slipped with ProLong gold antifade mounting medium (Life Technologies, Eugene, OR, USA). Two mice were excluded from data analysis due to considerable tissue damage.

### Electrophysiology

Sagittal (250 μm) sections of the NAcSh and NAcC were used for morphine challenge studies and acute morphine, with coronal slices (300 μm) used in morphine challenge and withdrawal studies as previously described (Hearing et al. 2016). Slices were collected in in a high sucrose solution as previously described (Hearing et al. 2013) and allowed to recover for at least 45-60 min in ACSF solution saturated with 95% O_2_/5% CO_2_ containing (in mM) 119 NaCl, 2.5 KCl, 1.0 NaH_2_PO_4_, 1.3 MgSO_4_, 2.5 CaCl_2_, 26.2 NaHCO_3_ and 11 glucose. Electrophysiological recordings assessing miniature excitatory postsynaptic currents (mEPSCs) were performed in the presence of picrotoxin (100μM) and lidocaine (700μM) to block GABAergic neurotransmission and sodium-dependent action potentials, respectively, as previously described (Hearing et al. 2016; Jedynak et al. 2016). The majority of NAcSh recordings were from the medial portion with an equal blend along the rostro-caudal axis. Cells were visualized using infrared-differential contrast (IR-DIC) microscopy, and medium spiny neurons (MSNs) were identified by cell subtype-specific fluorophore (tdTomato or EGFP) in combination with capacitance (>50 pF). Using a Sutter Integrated Patch Amplifier (Sutter Instruments, Novato, CA, USA) and/or Axon Instruments Multiclamp 700B (Molecular Devices, Sunnyvale, CA, USA), MSNs were voltage-clamped at −72 mV using electrodes (2.5 − 4 MΩ) with a cesium-methyl sulfonate based internal solution containing (in mM) 120 CsMeSO4, 15 CsCl, 10 TEA-Cl, 8 NaCl, 10 HEPES, 5 EGTA, 0.1 spermine, 5 QX-314, 4 ATP-Mg, and 0.3 GTP-Na. Data were filtered at 1-2 kHz and digitized at 20 kHz via custom Igor Pro software (Wavemetrics, Lake Oswego, OR, USA) or Clampex 10.7 software (Molecular Devices, Sunnyvale, CA, USA). Series (10-40 MΩ) and input resistance were monitored using a depolarizing step (5 mV, 100 ms). Neurons with a holding current below −150pA were excluded from analysis. Data collection and analysis were performed as previously described (Hearing et al. 2016; Kourrich et al. 2007).

Notably, independent samples t-tests between mEPSC metrics from morphine challenge study Sal-Sal mice recorded using sagittal (Univ of Minn) and coronal sections (Marquette Univ) were performed. We observed no impact of slice orientation/recording location on mEPSC metrics in NAcSh D1-MSNs (Amp t_(20)_=0.4359, p=0.67; Freq t_(20)_=0.5078, p=0.62), NAcSh D2-MSNs (Amp t_(11)_=0.8798, p=0.40; Freq t_(12)_=1.937, p=0.08), NAcC D1-MSNs (Amp t_(18)_=0.9552, p=0.35, Freq t_(20)_=0.5279, p=0.60), or NAcC D2-MSNs (Amp t_(16)_=1.86, p=0.08, Freq t_(16)_=1.88, p=0.08).

### Drugs

Picrotoxin and lidocaine were purchased from Sigma Aldrich (St. Louis, MO, USA). Morphine was purchased from the Boynton Pharmacy (University of Minnesota, Minneapolis, MN, USA) or Froedtert Hospital Pharmacy (Medical College of Wisconsin, Milwaukee, WI, USA).

### Statistical Analysis and Data Presentation

mEPSCs were analyzed with independent samples t-tests or two-way ANOVAs using SigmaPlot (Systat Software, San Jose, CA, USA) or Graph Pad Prism (GraphPad Software, Inc., La Jolla, CA, USA). Appropriate post hoc analyses were used for pairwise comparisons as indicated. The threshold for statistical significance in all cases was p<0.05. Electrophysiology data is represented utilizing standard column graphs displaying mean +/− SEM, with adjacent scatter plots of individual data points. Sample size in experiments is presented as *n* and *N*, where *n* is the number of cells and *N* is the number of mice.

## RESULTS

### Bi-directional changes in AMPA receptor synaptic transmission in the NAcSh and NAcC

We recently demonstrated that prolonged withdrawal from repeated non-contingent morphine increases excitatory drive at NAcSh D1-MSNs, while reducing drive at D2-MSNs (Hearing et al. 2016). Our previous work shows that re-exposure to relapse-inducing stimuli (i.e., discrete cues, drug, stress) following abstinence or extinction promotes a transient reduction in synaptic AMPAR signaling in unidentified MSNs that drives cocaine-induced reinstatement of place preference (Benneyworth et al. in press; Brebner et al. 2005; Ebner et al. 2018; Famous et al. 2008; Ingebretson et al. 2018; Jedynak et al. 2016; Kourrich et al. 2007; Rothwell et al. 2011; Schmidt et al. 2015; Schmidt et al. 2013; Thomas et al. 2001). While the cell-type selectivity of this plasticity remains unclear, previous reports indicate that this reduction occurs in D2-MSNs (Ortinski et al. 2015). Initial studies sought to determine whether re-exposure to opioids promotes a similar bi-modal shift in synaptic strength using *ex vivo* recordings of miniature excitatory postsynaptic currents (mEPSCs) – a direct measure of synaptic AMPAR function – 24 hrs following a post abstinence (10-14 d) challenge injection. To identify effects of acute morphine exposure based on previous drug experience and ensure effects of challenge injection are specific for drug re-exposure, this experiment contained 4 experimental groups. Two groups received 5 daily injections of saline and later received a saline or morphine priming injection (Sal-Mor, Mor-Mor), and two groups receiving 5 injections of morphine followed by a saline or morphine challenge (Mor-Sal, Mor-Mor), with recordings performed 24 hrs following the challenge injection. This approach permitted us to distinguish effects of repeated morphine and morphine re-exposure.

Two-way ANOVAs were performed on mEPSC amplitude and frequency with daily drug and challenge exposures as between subjects factors; Tukey *post hoc* analyses were performed when appropriate. In NAcSh D1-MSNs, mEPSC amplitude and frequency was elevated in Mor-Sal mice [amplitude: 14.85±0.83), frequency: (6.63±0.66)] compared to Sal-Sal controls [amplitude: (11.24±0.65), frequency: (5.99±1.01)], and that a morphine challenge (Mor-Mor) returned amplitude and frequency to control levels [amplitude: (11.27±0.35), frequency: (3.66±0.55)] (amplitude: interaction, F_(1,65)_=4.288, p<0.05); frequency: interaction, F_(1,65)_=14.08, p<0.001) (Figure 1B). While acute morphine exposure produced a trend toward increased mEPSC frequency compared to Sal-Sal mice, these effects were not significant. In contrast to D1-MSNs, no significant main effects or interactions were observed for D2-MSN mEPSCs amplitude (daily, F_(1,44)_=0.3648; challenge, F_(1,44)_=2.197, p=0.15; interaction, F_(1,44)_=0.2447) or frequency (daily, F_(1,44)_=0.0204; challenge, F_(1,44)_=1.468; interaction, F_(1,44)_=2.646, p=0.11) (Figure 1C). These data show that acute morphine exposure does not promote lasting alterations in glutamate plasticity in the NAcSh and that morphine re-exposure promotes a bimodal effect on AMPAR signaling akin to that observed following re-exposure to psychostimulants and associated stimuli (Benneyworth et al. ; Brebner et al. 2005; Ebner et al. 2018; Famous et al. 2008; Ingebretson et al. 2018; Jedynak et al. 2016; Kourrich et al. 2007; Rothwell et al. 2011; Schmidt et al. 2015; Schmidt et al. 2013; Thomas et al. 2001), however these effects may be confined to D1-MSNs unlike cocaine (Ortinski et al. 2015).

**Figure 1.**
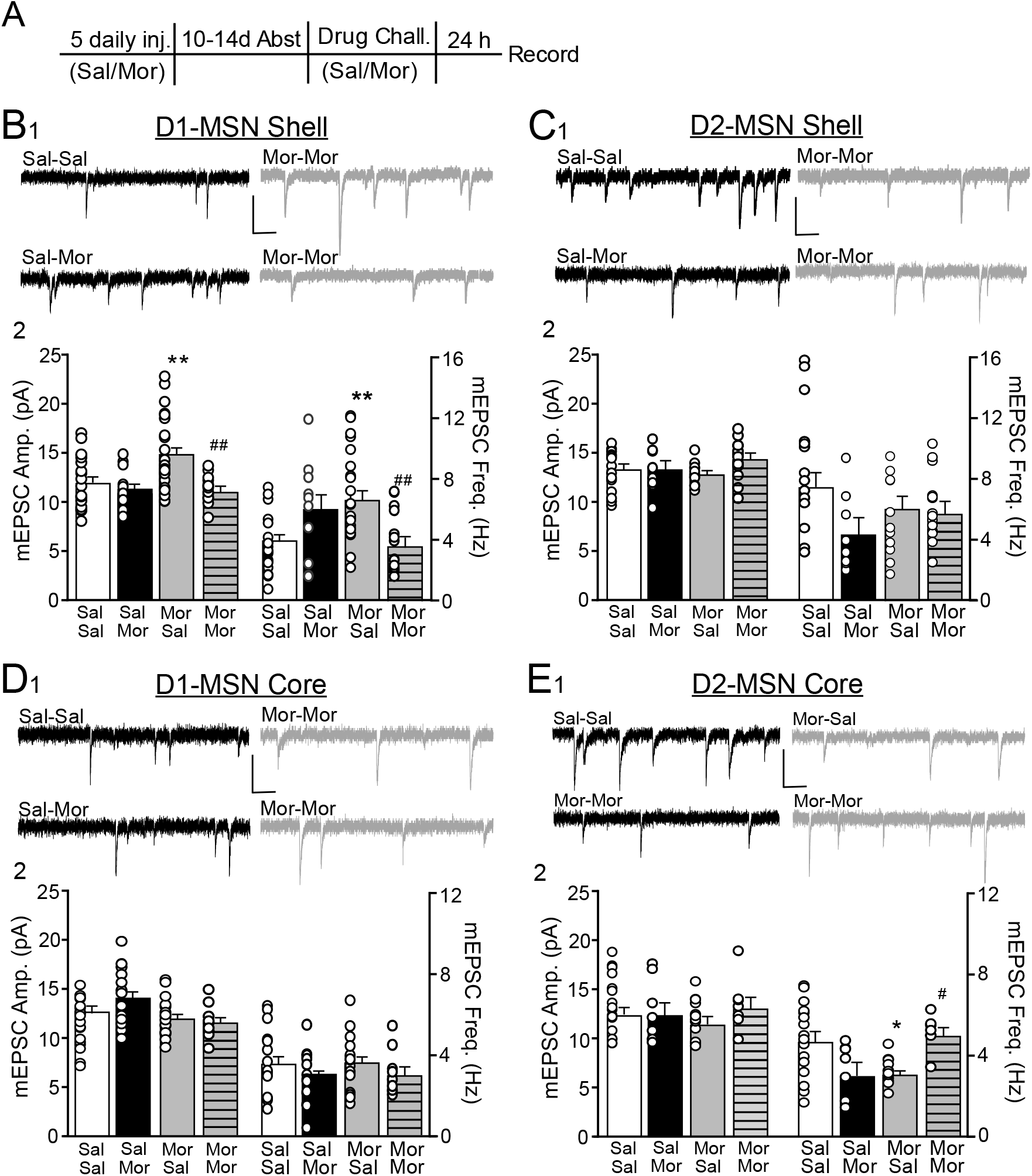
Cell- and region-specific effect of post-abstinence morphine exposure on nucleus accumbens AMPA receptor transmission. **A** Experimental timeline including 5 days of saline or morphine (10 mg/kg; i.p.), a 10-14 day withdrawal period, and challenge injection of saline or morphine. Electrophysiological recordings performed in NAcSh or NAcC D1- or D2-MSNs 24 hours post challenge injection in either coronal or sagittal slices (no significant difference was observed – see methods). **B** (1) Representative miniature excitatory postsynaptic current (mEPSC) traces and (2) mean mEPSC amplitude (left) and frequency (right) in NAcSh D1-MSNs from saline + saline challenge (Sal-Sal, white; n=20/N=11), saline + morphine challenge (Sal-Mor, black; n=11/N=6), morphine + saline challenge (Mor-Sal, gray; n=22/N=10) and morphine + morphine challenge (Mor-Mor, striped gray; n=15/N=5). **C** (1) Representative traces and (2) mean mEPSCs in NAcSh D2-MSNs [Sal-Sal (n=15/N=8),Sal-Mor (n=8/N=5), Mor-Sal (n=12/N=8), Mor-Mor (n=13/N=7)]. **D** (1) Representative traces and (2) mean mEPSCs in NAcC D1-MSNs [Sal-Sal (n=19/N=11), Sal-Mor (n=12/N=7), Mor-Sal (n=16/N=10), Mor-Mor (n=9/N=7). **E** (1) Representative traces and (2) mean mEPSCs in NAcC D2-MSNs [Sal-Sal (n=17/N=11), Sal-Mor (n=7/N=6), Mor-Sal (n=13/N=10), Mor-Mor (n=6/N=5) Scale bar = 20 pA/100 ms; Tukey post hoc: *p<0.05, **p<0.01 vs Sal-Sal, #p<0.05, ##p<0.01 vs Mor-Mor.

In the NAcC, mEPSC amplitude in D1-MSNs from Sal-Sal (12.47±0.59) and Sal-Mor (14.13±0.97) were significantly higher compared to Mor-Sal (11.86±0.54) and Mor-Mor (11.53±0.66) (main effect of daily, F_(1,57)_=6.447, p<0.05), whereas no effects were observed for D1-MSN mEPSC frequency (daily, F_(1,55)_=0.0696; challenge, F_(1,55)_=2.483, p=0.12; interaction, F_(1,55)_=0.0693). Examination of D2-MSNs showed that mEPSC frequency was reduced in Mor-Sal (3.06±0.19) compared to Sal-Sal (4.77±0.48), and that a morphine challenge (Mor-Mor) returned mEPSC frequency to Sal-Sal control levels (5.44±0.22) (interaction: F_(1,40)_=13,96, p<0.001) (Figure 1C). No effects were observed for NAcC D2-MSN mEPSC amplitude (daily, F_(1,43)_=0.0178; challenge, F_(1,43)_=0.682; interaction, F_(1,43)_=0.6136). These data align with our previous findings that repeated morphine reduces excitatory drive at NAcC D2-MSNs without altering transmission at D1-MSNs, and this plasticity is bimodal in nature similar to plasticity at D1-MSNs in the NAcSh,.

### Effects of acute morphine on nucleus accumbens cell-specific AMPAR signaling

Despite ample evidence indicating that glutamate transmission at NAc MSNs contributes to the rewarding effects of opioids (Dworkin et al. 1988; Graziane et al. 2016; Hearing et al. 2016; Olds 1982; Vekovischeva et al. 2001), our data show that a single morphine injection has no significant impact on AMPAR mEPSC signaling in the NAcSh or NAcC 24 hrs after exposure. Previous data has shown that psychostimulant-induced plasticity in the NAc can be induced following a single exposure but requires time to develop (Kourrich et al. 2007; Terrier et al. 2016). As there are no known data on the immediate effects of acute *in vivo* opioid exposure on AMPAR transmission in the NAc (Chartoff and Connery 2014), we next investigated the impact of acute opioid exposure on AMPAR synaptic plasticity in the NAc approximately 3 hrs following an injection of saline or morphine in D1- and D2-MSNs of the NAcSh and NAcC. This timepoint was choicen to isolate reward-versus withdrawal-related effects of morphine as it aligns with elevations in drug-induced motor activity (not shown) and resides within the approximate 5-hour half-life of subcutaneous morphine administration (Hipps et al. 1976).

In the NAcSh, mEPSC amplitude and frequency were significantly elevated in D1-MSNs of morphine-treated animals compared to saline controls (Figure 2B) [amplitude: Sal (11.40±0.64), Mor (16.25±1.28), t_(16)_=3.07, p<0.01); frequency: Sal (5.02±0.46), Mor (9.90±1.5), t_(16)_=2.79, p<0.05]. Alternatively, acute morphine increased mEPSC frequency but not amplitude at D2-MSNs (Figure 2C) ([amplitude: Sal (15.2±0.19), Mor (14.2±0.81), t_(6)_=1.22, p=0.27; frequency: Sal (5.95±0.53), Mor (10.73±1.05), t_(6)_=4.06, p<0.01].

**Figure 2.**
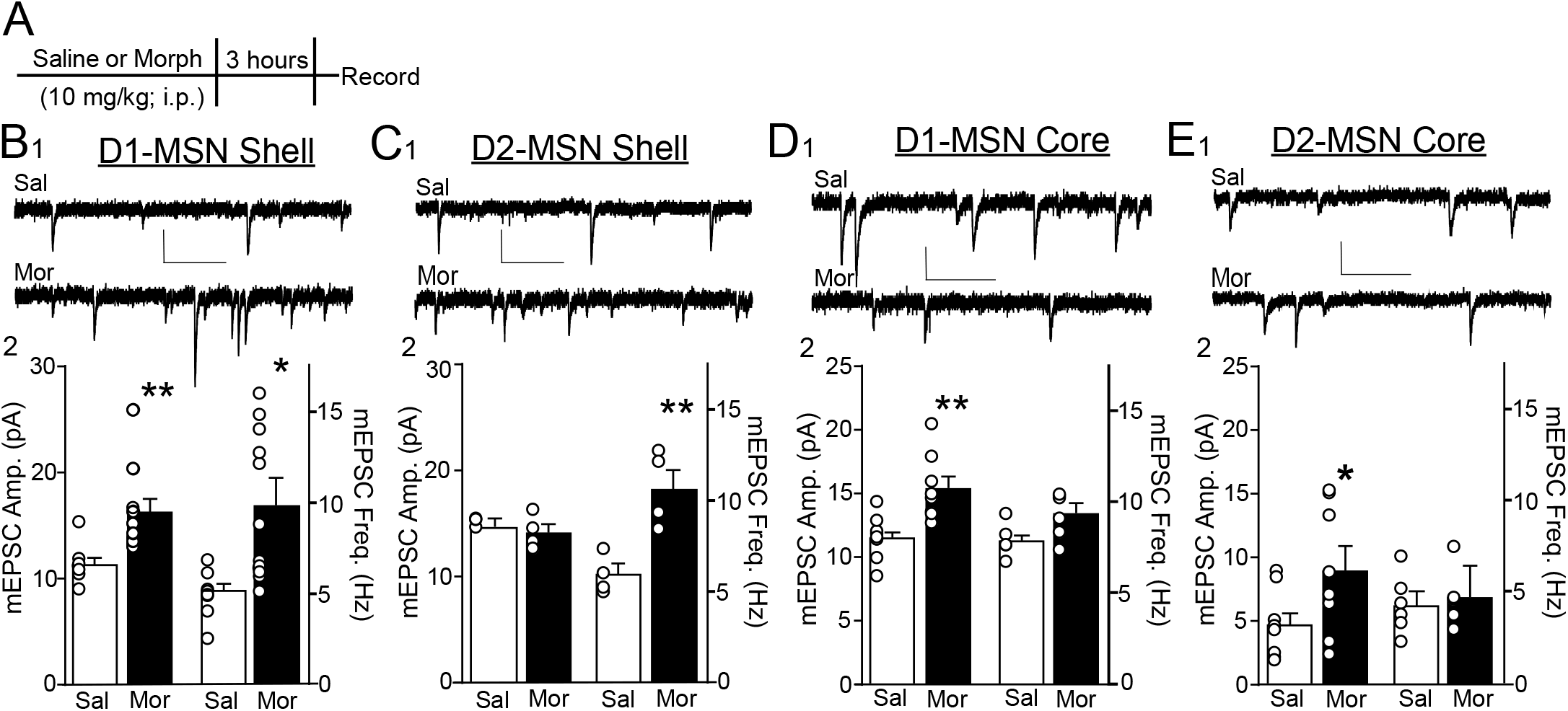
Cell-specific effects of acute morphine on nucleus accumbens AMPA receptor transmission. **A** Experimental timeline including an acute injection of saline or morphine 10 mg/kg; i.p.)and electrophysiological recordings performed 3-4 hours post injection. Recordings were performed in sagittal slices containing the NAc shell or core. **B** (1) Representative AMPAR mEPSC traces and (2) mean mEPSC amplitude (left) and frequency (right) in D1-MSNs from saline (Sal, white; n=8, N=5 and morphine (Mor, black; n=10/N=5) injected mice. **C** (1) Representative traces and (2) mean mEPSCs in NAcSh D2-MSNs (Sal: n=4/N=4; Mor: n=4/N=4). **D** (1) Representative traces and (2) mean mEPSCs in NAcC D1-MSNs (Sal n=9/N=6; Mor: n=8/N=7). **E** (1) Representative mEPSC traces and (2) mean mEPSCs in NAcC D2-MSNs (Sal: n=5, N=5; Mor: n=5/N=4). *p<0.5, **p<0.01 vs Sal. Scale bars = 20 pA/100 ms.

Similar to the NAcSh, acute morphine increased mEPSC amplitude and frequency at D1-MSNs in the NAcC (figure 2D) [amplitude: Sal (11.54±0.55), Mor (15.43±0.92), t_(15)_=3.72, p<0.01; frequency: Sal (3.24±0.59), Mor (6.23±1.3), t_(15)_=2.23, p<0.05]. However, neither amplitude or frequency of mEPSCs was altered by acute morphine at NAcC D2-MSNs (Figure 2E) [amplitude: Sal (11.33±0.62), Mor (13.32±0.73), t_(9)_=2.022, p=0.074; frequency: Sal (4.36±0.66), Mor (4.74±0.77), t_(9)_=0.382, p=0.71]. Taken together, these data indicate that acute morphine promotes a global augmentation of excitatory drive at D1-MSNs, whereas alterations in glutamate transmission at D2-MSNs are specific to the NAcSh, and that these adaptations do not persist or require a period of abstinence to manifest.

### Spontaneous withdrawal enhances AMPAR signaling at D2-MSNs

In addition to our lack of knowledge about the effects of acute *in vivo* opioids on NAc synaptic transmission, few studies to date have explored the impact of withdrawal-inducing morphine administration on glutamate transmission in a NAc sub region and MSN subtype specific manner. A single injection of morphine is sufficient for precipitated withdrawal symptoms 24 hrs after exposure (Rothwell et al. 2012). Further, prior studies indicate that NAcSh D2-MSNs potently regulates somatic withdrawal symptoms (Harris and Aston-Jones 1994; Russell et al. 2016; Zhu et al. 2016). This is significant given our observed plasticity 3 hrs, but not 24 hrs after a single injection of morphine. Surprisingly, no studies to date have examined cell- or region-specific NAc plasticity associated with *spontaneous* somatic withdrawal despite a purported role in relapse behavior. In order to assess adaptations associated with spontaneous withdrawal and determine whether plasticity is uniquely different following a similar withdrawal period (24 hrs) but a different regimen, we administered an escalating dose of morphine shown to produce dependence as measured by spontaneous withdrawal symptoms (Papaleo and Contarino 2006) and recorded metrics of spontaneous withdrawal 2 hrs prior to preparation for acute slice electrophysiology. Escalating doses of morphine significantly increased the number of jumps (t_(17)_=2.217, p<0.05), tremors (t_(17)_=2.89, p<0.05), wet dog shakes (t_(17)_=2.503, p<0.05), and piloerection (t_(17)_=5.037, p<0.001) (table 1). No significant effect of morphine was observed on grooming behavior (t_(17)_=0.729, p=0.48) or paw flutters (t_(17)_=0.527, p=0.61) (table 1). More specifically, morphine significantly increased the total withdrawal score (figure 3) [Sal (8.19±1.57), Mor (20.06±3.14, t_(17)_=2.741, p<0.05)], and decreased distance traveled in a novel context [Sal (85.73±8.27), Mor (38.76±2.13, t_(17)_=6.906, p<0.0001)]. Notably, in a separate cohort of mice, we examined whether somatic withdrawal symptoms persisted at 10-14 d post drug exposure – a timepoint examined in drug challenge studies. Two-way day-by-drug ANOVA with day a repeated measure and drug exposure as a between subjects factor reveal significant main effects of day (F_(1,13)_=10.52, p<0.01), drug (F_(1,13)_=4.481, p=0.05), and a significant day-by-drug interaction (F_(1,13)_=12.08, p<0.01). Bonferroni post hoc analyses showed no significant impact of day on saline-exposed mice (p=0.99) but a significant decrease in withdrawal score after 14-day abstinence in morphine-exposed mice (p<0.001).

**Figure 3.**
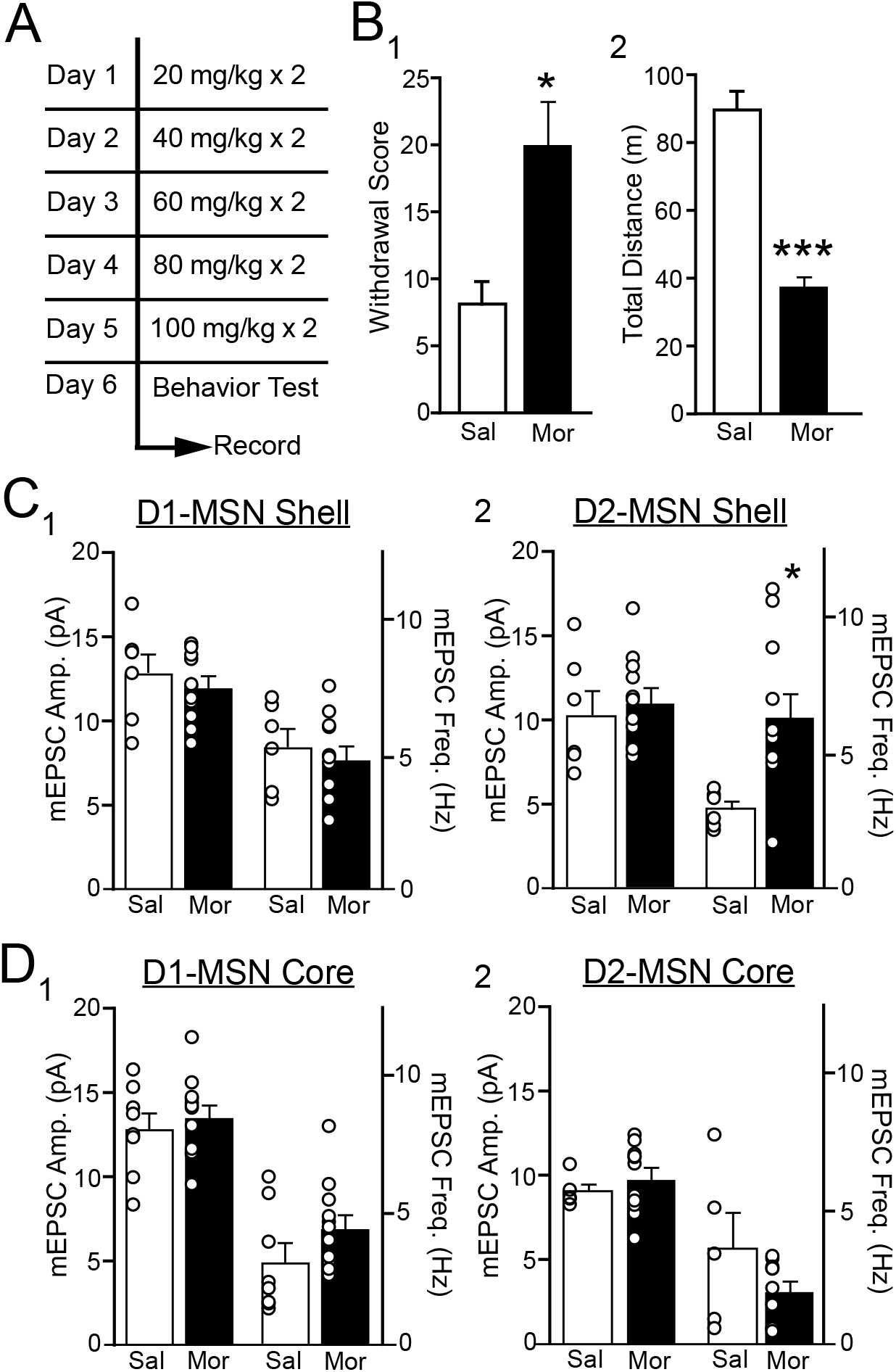
Cell-specific effects of morphine-induced spontaneous acute withdrawal on nucleus accumbens AMPA receptor transmission. **A** Experimental timeline including five twice-daily injections of escalating morphine administration (i.p.) and 24 hours post final injection behavior assessment. Recordings were performed in coronal slices approximately 2 hours following behavior. **B** Mean (1) global withdrawal scores and (2) locomotor activity 24 hours following repeated saline (Sal, white; N=7) or morphine (Mor, black; N=12) injections. **C** Mean mEPSC amplitude (left) and frequency (right) in NAcSh (1) D1-MSNs [Sal (n=6/N=5), Mor n=11/N=7)] and (2) D2-MSNs [Sal (n=6/N=5), Mor (n=12/N=8)]. **D** Mean mEPSC amplitude and frequency in NAcC (1) D1-MSNs [Sal (n=8/N=7), Mor (n=12/N=9)] and D2-MSNs [Sal (n=5/N=4), Mor (n=11/N=8). *p<0.05, ***p<0.001 vs Sal.

To identify plasticity that parallel withdrawal symptoms, we measured mEPSCs at D1- and D2-MSNs in the NAcC and NAcSh subregions 24 hrs following the final injection of morphine and 2 hrs post behavior assessment. In the NAcSh, neither the amplitude or frequency of mEPSCs were altered in D1-MSNs (Figure 3C) [amplitude: Sal (12.75±1.23), Mor (12.01±0.61), t_(15)_=0.612, p=0.55; frequency: Sal (5.58±0.72), Mor (4.84±0.50), t_(15)_=0.862, p=0.40]. Conversely, a significant increase in frequency but not amplitude was observed in D2-MSNs (Figure 3C): [amplitude: Sal (10.39±1.41), Mor (11.12±0.81), t_(15)_=0.487, p=0.63; frequency: Sal (2.94±0.27), Mor (6.40±0.86), t_(15)_=2.876, p<0.05]. In the NAcC, morphine had no effect on mEPSC amplitude or frequency in D1-MSNs (Figure 3D) [amplitude: Sal (12.76±0.94), Mor (13.49±0.67), t_(18)_=0.655, p=0.52; frequency: Sal (3.01±0.68), Mor (4.25±0.47), t_(18)_=1.558, p=0.14] or D2-MSNs (Figure 3D) [amplitude: Sal (9.01±0.42), Mor (9.80±0.60), t_(14)_=0.832, p=0.42; frequency: Sal (3.52±1.34), Mor (1.9±0.33), t_(14)_=1.536, p=0.15]. Taken together, these findings suggest that, similar to naloxone-precipitated withdrawal (Zhu et al. 2016), spontaneous somatic withdrawal selectively increases excitatory drive at D2-MSNs in the NAcSh and that these effects are more enduring than previously known. Further, these data support the notion that adaptations 10-14 d following a less robust morphine regimen (5 × 10 mg/kg) or lack thereof 24 hrs following acute exposure is not associated with enduring somatic withdrawal, and that the lack of plasticity observed 24 hrs following an acute injection is distinctly different from withdrawal associated plasticity at a similar timepoint.

### Inhibiting endocytosis of AMPA receptors in the NAcSh enhances morphine-primed reinstatement of place preference

Inhibiting NAc GluA2-containing AMPAR trafficking disrupts amphetamine-induced sensitization and attenuates cocaine-induced reinstatement of cocaine-seeking and place preference (Benneyworth et al. in press; Brebner et al. 2005; Famous et al. 2008), suggesting that reductions in AMPAR signaling may reflect transient plasticity that triggers relapse-related behavior. Therefore, we examined whether morphine-induced depotentiation of AMPAR-signaling involves receptor endocytosis and if this plasticity is causally involved in reinstatement of reward behavior. As plasticity was largely confined to the NAcSh, we focused our efforts for this experiment in this sub-region. Using an approach previously shown to produce AMPAR plasticity and conditioned place preference (Hearing et al. 2016), all mice were initially conditioned with morphine (5 mg/kg). Mice were subsequently divided into three experimental groups, one group infused with the active peptide (Tat-GluA2_3Y_) and receiving a saline priming injection (Mor/Tat(+)/Sal), a second group receiving the active peptide and a priming injection of morphine (Mor/Tat(+)/Mor), and a third infused with the inactive (Tat-GluA2_3S_) isoform and receiving a morphine prime (Mor/Tat(-)/Mor).

A Two-way ANOVA with test day as a repeated measure and treatment group as a between subjects factor revealed a significant day-by-treatment interaction (F_(6,135)_=2.84, p<0.05). Post hoc pairwise multiple comparisons showed that all three groups exhibited significant place preference compared to pre-test preference levels, and that preference did not significantly differ across all three groups during pre-test, preference, or extinction. For reinstatement testing, mice infused with the inactive Tat-GluA2_3S_ peptide receiving a morphine prime displayed a significant increase in preference compared to Tat-GluA23Y infused mice injected with saline (Figure 4B_1_) [Mor/Tat(+)/Sal (45.52±83.7), Mor/Tat(-)/Mor (449±79.05); p<0.001], while morphine primed mice infused with the active peptide (Mor/Tat(+)/Mor: 666±87.8) showed a significant increase in preference reinstatement compared to both groups (p<0.05 vs. Mor/Tat(-)/Mor; p<0.001 vs. Mor/Tat(+)/Sal], indicating that blockade of AMPAR endocytosis enhanced reinstatement of place preference. Approximately 30-90 min following testing, *ex vivo* analysis of mEPSCs (Figure 4B_2_) showed that mEPSC amplitude at D1-MSNs was significantly greater in mice infused with active Tat-GluA23Y compared to the inactive Tat-GluA23S [Mor/Tat(-)/Mor 16.76±1.87, Mor/Tat(+)/Mor 11.24±0.32, t_(6)_=2.911, p<0.05]. No significant effect of Tat peptide on mEPSC frequency was observed [Mor/Tat(-)/Mor (3.25±0.48), Mor/Tat(+)/Mor (4.35±0.95), t_(6)_=1.032, p=0.34]. Collectively, these data indicate that re-exposure to morphine drives endocytosis of AMPA receptors at D1-MSNs unlike cocaine (Benneyworth et al. in press), and preventing such endocytosis exacerbates reinstatement of morphine-induced conditioned place preference.

**Figure 4.**
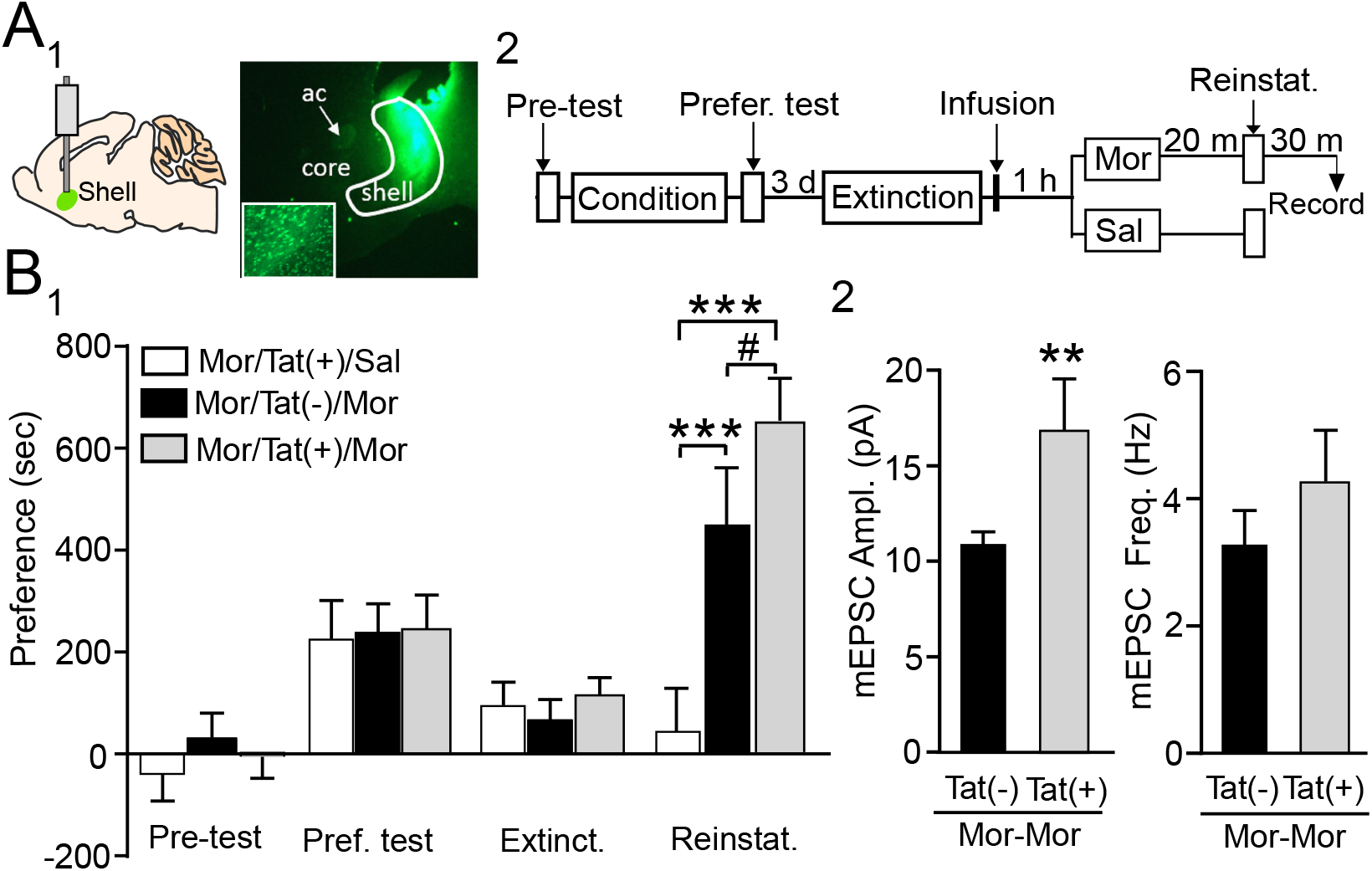
Inhibiting AMPA receptor endocytosis in nucleus accumbens shell increases morphine-induced reinstatement of conditioned place preference. **A** (1) Representative Tat-peptide expression targeted to the NAc shell and (2) experimental timeline of conditioned place preference study including behavioral testing. Electrophysiological recordings were performed in Tat-expressing D1-MSNs fro sagittal slices approximately 30 min post reinstatement test. All mice were conditioned with morphine. Following infusion with the active (Tat+) or inactive (Tat-) peptide, mice received a challenge injection of saline (Mor/Tat+/Sal, white), or morphine (Mor/Tat(-)/Mor, black; Mor/Tat(+)/Mor, gray), followed by a test of preference reinstatement 20 min later. **B** (1) Preference scores across test days in mice and(2) mEPSC amplitude (left) and frequency (right) from D1-MSNs expressing fluorescence following testing in a subset of mice receiving morphine injections with the active or inactive peptide. #p<0.05 Mor/Tat(-)/Mor vs. Mor/Tat(+)/Mor, ***p<0.001 vs. Mor/Tat+/Sal.

## DISCUSSION

Here we identify temporal- and region-specific changes in AMPAR signaling within discrete subpopulations of NAc MSNs associated with opioid reward, withdrawal, and relapse-like behavior. In agreement with our previous findings, we showed that protracted withdrawal from repeated morphine is associated with increases in synaptic drive at D1- and decreases at D2-MSNs predominantly in the NAcSh (Hearing et al. 2016). Similar to amphetamine and cocaine (Benneyworth et al. in press; Jedynak et al. 2016; Kourrich et al. 2007), re-exposure to morphine produced bimodal plasticity, however, unlike cocaine, re-exposure to morphine triggered a reduction in drive at D1- and increased drive at D2-MSNs (Figure 1; Ortinski et al., 2015). In contrast, acute morphine produced a transient increase in AMPAR-mediated neurotransmission at D1-MSNs in the NAcC *and* NAcSh (Figure 2) whereas spontaneous withdrawal aligned with enhanced excitatory drive at D2-MSNs selectively in the NAcSh (Figure 3). Unlike our previous findings with cocaine, blocking the underlying AMPAR endocytosis augmented rather than inhibited reinstatement of place preference following extinction training (Figure 4), suggesting that similar forms of plasticity and post-drug experience may have distinct behavioral consequences across drug classes.

### Acute morphine plasticity

Despite evidence that opioids acutely reduce glutamate release in the NAc (Martin et al. 1997; Sepulveda et al. 1998), relatively little is known regarding the role of NAc postsynaptic AMPAR and NMDAR signaling in the acute rewarding effects of opioids. Biochemical data have shown that expression of AMPARs is decreased in the NAcC 3 days following an acute morphine exposure (Jacobs et al. 2005) and that GluA1 surface expression is reduced in combined NAc tissue 24 hrs following acute exposure (Herrold et al. 2013). In the present study, we found that AMPAR-mediated mEPSC amplitude and frequency was elevated at D1-MSNs in the NAcC and NAcSh 3 hrs following an acute injection of morphine. Unlike previous studies, this time course aligns with residual motor activity following drug exposure as well as the acute rewarding effects of morphine rather than withdrawal (Rothwell et al. 2012). The NAcC plays a key role in initializing reward-related motor activity (Sesack and Grace 2010; Shiflett and Balleine 2011; Voorn et al. 2004) and the NAcSh in opioid-related reward and reinforcement learning (Graziane et al. 2016; Hearing et al. 2016; Heimer et al. 1997; Sesack and Grace 2010). Moreover, recent findings have shown that activation of NAc D1-MSN circuits promotes reward and addiction-related behavior (Dobi et al. 2011; Graziane et al. 2016; Hearing et al. 2018; Hearing et al. 2016; Kim et al. 2011; Lobo et al. 2010; Ortinski et al. 2015; Ortinski et al. 2012; Pascoli et al. 2011; Smith et al. 2013; Suska et al. 2013). Thus, increased signaling at D1-MSNs in the NAcSh and NAcC likely contribute to the rewarding and psychomotor effects of acute morphine, respectively. This is reflected in enhanced drug-induced behavioral output observed after repeated and acute morphine (Hearing et al. 2016); behavior blunted by depotentiation of postsynaptic AMPAR signaling.

It remains unclear whether elevations in AMPAR transmission at D1-MSNs in the NAcSh and NAcC reflect pre- or postsynaptic events. Accordingly, observed increases in mEPSC frequency may be attributed to increased receptor (or synapse) number rather than glutamate release probability (Graziane et al. 2016; Kerchner and Nicoll 2008) given the dampening effects of acute opioids on glutamate release in the nucleus accumbens (Sepulveda et al. 2004). Recent work has shown that a single cocaine exposure upregulates NAcSh GluA2-lacking AMPAR signaling at NAcSh D1R-MSN synapses – an adaptation observed 7, but not 1 day following drug exposure (Terrier et al. 2016). Although an in-depth comparison of cocaine-induced changes at the acute post exposure period used here is lacking, acute morphine adaptations in the NAcSh also appear to require a period of withdrawal, as they were not observed 24 hrs following exposure. Alternatively, while mEPSC amplitude and frequency and GluA1 surface expression are elevated in pooled MSNs and tissue punches during early withdrawal from acute amphetamine and repeated cocaine, no changes in mEPSCs were observed 24 hrs following acute morphine in the present study. Although the reason for this distinction is unclear, it may reflect a higher prevalence of mu opioid receptors in the NAcSh compared to the NAcC (Svingos et al. 1997). Regardless, given increasing evidence that opioids and psychostimulants produce divergent neurophysiological and behavior effects, an important question moving forward will be to determine whether increased AMPAR signaling with acute morphine merely reflects a synaptic scaling event in response to reduced glutamate availability or if this plasticity persists.

### Withdrawal-related AMPAR plasticity

In addition to reward, increased AMPAR signaling in the NAcSh has also been attributed to aversive effects of morphine withdrawal (Russell et al. 2016; Sepulveda et al. 2004). Indirect pharmacological evidence as well as direct measures of synaptic plasticity indicate that these adaptations may be confined to D2-MSNs in the NAc (Harris and Aston-Jones 1994; Russell et al. 2016; Zhu et al. 2016). In the present study we show for the first time that, similar to naloxone-precipitated withdrawal, spontaneous somatic withdrawal aligns with increased glutamate transmission selectively at D2-MSNs in the NAcSh. Surprisingly, a similar phenomenon was also observed immediately following acute morphine exposure in both instances, though effects were confined to changes in mEPSC frequency. This may reflect increases in quantal release from pooled inputs, as withdrawal-related aversion memories are associated with increased signaling at thalamus but not prefrontal cortex or amygdala inputs at D2-MSNs (Zhu et al. 2016), but not definitively excluding a potential change in AMPAR expression (Graziane et al. 2016; Kerchner and Nicoll 2008). Importantly, we also observed that acute morphine increased signaling at NAcSh and NAcC D1-MSNs, possibly offsetting of the negative affect of a single post-morphine exposure.

Although a single exposure to morphine does not evoke spontaneous withdrawal, naloxone-precipitated withdrawal is possible 24 hrs after a single injection of morphine (Rothwell et al. 2012). Thus, our findings appear to agree with conclusions drawn by Russel et al., (2016) in that upregulation of glutamate transmission (at D2-MSNs in the present study) may reflect plasticity that primes NAc circuits for subsequent activation upon withdrawal (Russell et al. 2016). Although unclear, the apparent discrepancy between observed reductions in GluA2-lacking AMPAR surface expression immediately following precipitated withdrawal (Russell et al. 2016) and lack of changes to amplitude in the present study may reflect distinctions in the time of observation (30 min vs. 24 hrs) or method of withdrawal (precipitated vs spontaneous). Alternatively, because mEPSCs likely reflect binding at receptors in the synapse, it is possible that reduced surface expression detected using biochemical measures (e.g., biotinylation) reflect sampling from synaptic *and* peri/extra-synaptic AMPARs that have been primed but not trafficked to or from the postsynaptic density.

### Region- and cell-specific bimodal AMPAR plasticity

Our previous findings show that 10-14 days after repeated morphine increases expression of GluA2-lacking AMPARs at pooled inputs to D1-MSNs while reducing excitatory drive at D2-MSNs in the NAcSh and NAcC (Hearing et al. 2016). In the present study, we also observed reductions in mEPSC frequency at NAcC D2-MSNs, but only a trend towards reduced signaling at NAcSh D2-MSNs. The prominence of plasticity in the NAcSh vs NAcC appears to contrast effects of repeated cocaine, but is consistent with findings following 10-14 d withdrawal from repeated amphetamine (Jedynak et al. 2016), however these effects have not been readily characterized in D1-vs D2-MSNs. In the current study, reductions in AMPAR signaling following morphine re-exposure were ostensibly isolated to D1-MSNs (Figure 1B). In turn, morphine treated mice infused with the active Tat peptide in the NAcSh exhibited increased mEPSC amplitudes compared to those infused with the inactive form (Figure 4B). Thus, morphine re-exposure likely triggers endocytosis of synaptic GluA2-containing AMPARs in NAcSh D1-MSNs. Further, as the Tat peptide inhibits activity-dependent rather than constitutive removal of synaptic GluA2-containing AMPARs, this endocytosis is more likely to reflect a rapid, LTD-like process than a slow and consistent removal of synaptic AMPARs over time (Ahmadian et al. 2004; Dong et al. 2015; Lee et al. 2002; Scholz et al. 2010; Wang et al. 2017; Yoon et al. 2009).

It should be noted that the precision of Tat injections in the current study was not specific with regards to the rostral-caudal and dorsal-ventral axis. This is significant as prior work has shown distinctions in how NAcSh cell subpopulations and AMPAR signaling along the dorsal-ventral and rostro-caudal gradient differentially regulate reward- and aversive-driven behavior (Reynolds and Berridge 2003). As our recordings were primarily, but not exclusively, focused within the dorsal-medial region with even distribution along the rostral-caudal axis, it will be important for future studies to examine anatomical distinctions when identifying causality between plasticity and behavior.

Our current findings indicate that re-exposure to morphine promotes of AMPAR endocytosis specifically at D1-MSN synapses previously potentiated during withdrawal from morphine, however it is possible that reductions in synaptic strength may be occurring at adjacent, rather than previously potentiated synapses – a possibility difficult to demonstrate definitively. Further, although it is impossible to exclude, it is unlikely that inclusion of extinction produced alternate plasticity in CPP studies compared to those observed in challenge experiments involving home cage abstinence, as our previous work showed identical cell-specific plasticity in mice following home cage abstinence and extinction. While the significance of this bimodal phenomenon is not yet clear, one intriguing possibility is that re-exposure to opioids may represent a temporary quelling of drug craving and in turn a trend towards returning to prior levels of D1-MSN excitation.

### Role of bidirectional plasticity in reinstatement

Our previous work showed that *in vivo* reversal of morphine-induced pathophysiology at NAcSh D1-MSNs with optogenetic stimulation or treatment with the antibiotic, ceftriaxone, blocked reinstatement of morphine-evoked place preference (Hearing et al. 2016). One straightforward interpretation of these findings is that a progressive enhancement of AMPAR signaling during withdrawal serves as a common mechanism for driving addiction-related behavior (Kalivas 2009; Kalivas and Hu 2006), and that reducing synaptic strength prior to drug re-exposure impairs drug-induced behavior. This contention is also supported by numerous studies showing that re-exposure to drug or drug-associated cues induces a rapid potentiation (i.e., release) of NAc glutamatergic signaling in cocaine, nicotine, or heroin withdrawn rats (Gipson et al. 2013a; Gipson et al. 2013b; Shen et al. 2011; Trantham-Davidson et al. 2012).

On the other hand, recent work by our group and others indicate that re-exposure to cocaine triggers a rapid reduction in synaptic strength in the NAc akin to LTD (Benneyworth et al. in press; Ebner et al. 2018; Ingebretson et al. 2018; Jedynak et al. 2016), and that this short-term plasticity is necessary and sufficient to reinstate place preference (Benneyworth et al. in press) – suggesting that *decreases* in excitatory drive onto NAc MSNs, particularly in the NAcSh, may promote reinstatement to drug seeking. Thus, our previously observed blockade of preference reinstatement may merely reflect an occlusion of short-term plasticity associated with morphine re-exposure and the ability to trigger behavior (Hearing et al. 2016; Pascoli et al. 2014; Pascoli et al. 2011). In the present study, blockade of AMPAR endocytosis augmented reinstatement of place preference, thus, unlike cocaine, increased expression of AMPAR during abstinence appears to be the primary driver of reinstatement, with internalization of AMPARs following morphine re-exposure perhaps reflecting a secondary synaptic scaling (Turrigiano 2008) in response to augmented glutamate release, but see (Siahposht-Khachaki et al. 2017). Regardless, these findings show that although two distinct drug classes can produce seemingly similar forms of plasticity, the behavioral consequences of this plasticity appear to be profoundly different.

### Conclusion

Though psychostimulants and opioids share rewarding properties that can lead to uncontrollable drug use and relapse vulnerability, opioids are addictive substances with the ability to induce chemical dependence from which relapse is driven by attempts to alleviate somatic and psychological withdrawal symptoms. By modeling dosing regimens in a preclinical setting, we sought to parallel acute, repeated, and dependence-inducing opioid consumption. Analysis of AMPAR signaling from each dosing revealed unique and overlapping neuroplasticity to excitatory signaling at NAcSh and NAcC MSNs. Future therapeutic interventions should take into consideration that drug-induced neuroplasticity shared across drug classes does not inherently share functional consequences at the level of the neural circuit or in terms of behavior. Thus, more thorough characterization of opioid-induced plasticity is needed to provide more efficient and effective therapies for opioid use disorder.

## Acknowledgements

The behavioral work done in part with the support by the Mouse Behavior Core at the University of Minnesota, which received funding from the National Institute for Neurological Disorders and Stroke (P30 NS062158). These studies were further supported by funding from the National Institute on Drug Abuse grant K99 DA038706 (to M.H.), R00DA038706 (M.H.), R00DA038706-04S1 (A.C.M), R01DA019666 (M.J.T.), K02DA035459 (M.J.T.) and T32 DA007234 (A.E.I.).

